# Cell surface crowding is a tunable biophysical barrier to cell-cell fusion

**DOI:** 10.1101/2024.12.12.628283

**Authors:** Daniel S.W. Lee, Liya F. Oster, Sungmin Son, Daniel A. Fletcher

**Affiliations:** Department of Bioengineering, University of California, Berkeley, CA 94720; Graduate Group in Biophysics, Berkeley, CA 94720; Division of Biological Systems and Engineering, Lawrence Berkeley National Laboratory, Berkeley, CA 94720; Chan Zuckerberg Biohub, San Francisco, CA 94158

## Abstract

Cell-cell fusion is fundamental to developmental processes such as muscle formation, as well as to viral infections that cause pathological syncytia. An essential step in fusion is close membrane apposition, but cell membranes are crowded with proteins, glycoproteins, and glycolipids, all of which must be cleared before a fusion pore can be nucleated. Here, we find that cell surface crowding drastically reduces fusogenicity in multiple systems, independent of the method for driving fusion. We estimate that cell surface crowding presents an energetic barrier to membrane apposition on the scale of ∼100*k*_*B*_*T*, greater than that of bare membrane fusion. We show that increasing cell surface crowding reduces fusion efficiency of PEG-mediated and fusogen-mediated cell-cell fusion, as well as synthetic membranes under force. Interestingly, we find that differentiating myoblasts naturally decrease cell surface crowding prior to fusion. Cell surface crowding presents an underappreciated biophysical barrier that may be tuned developmentally and could be targeted externally to control tissue-specific cell-cell fusion.

## Introduction

Cell-cell fusion is critical for the formation of specialized multinucleated cells in varied biological systems. In vertebrates ranging from lamprey eels to humans, fusion of myoblasts is required for development and maintenance of skeletal muscle fibers^1^. Similarly, multinucleated osteoclasts are crucial for bone resorption^2^. In mammals, fusion of mononucleated trophoblasts into the multinucleated placental syncytiotrophoblast provides a protective environment for embryo development^3^. Some viruses are also known to cause fusion of host cells. Fusion of respiratory epithelial cells by the SARS-CoV-2 virus can indicate severity of an infection^4^, and members of the orthoreovirus family cause fusion in their infected hosts, aiding in viral spread^5,6^. Each of these physiologically diverse examples of cell-cell fusion is driven by distinct membrane fusion proteins, called fusogens. Typically broken into three major classes based on their different structures, fusogens function through a wide variety of mechanisms, including disrupting bilayer packing to facilitate lipid mixing of apposed bilayers, inducing bilayer curvature favorable for hemifusion stalk or pore formation, and recruiting necessary accessory proteins to the fusion synapse.^7,8,9,10^

Despite structural and mechanistic differences in fusogens, lipid bilayers that fuse are believed to follow a qualitatively similar pathway to overcome the considerable energetic barrier to fusion, which for bare lipid bilayers has been estimated to be approximately 40*k*_*B*_*T*^11,12^. First, the membranes must come into close apposition. They then undergo hemifusion, in which the outer membranes fuse, forming a stalk, followed by full fusion, in which the hemifusion stalk opens into a fusion pore^11^. While this pathway is consistent with in vitro membrane fusion experiments and simulations^13,14^, it leaves out aspects of real cell-cell interactions that could significantly affect this fusion landscape and resulting fusion efficiency. In particular, the first step, during which membranes are brought into close apposition, requires exclusion of surface molecules and solvent from the intermembrane space (Fig. 1A). This raises the question of whether cell surface crowding is a significant barrier that could limit cell-cell fusion.

**Fig. 1.**
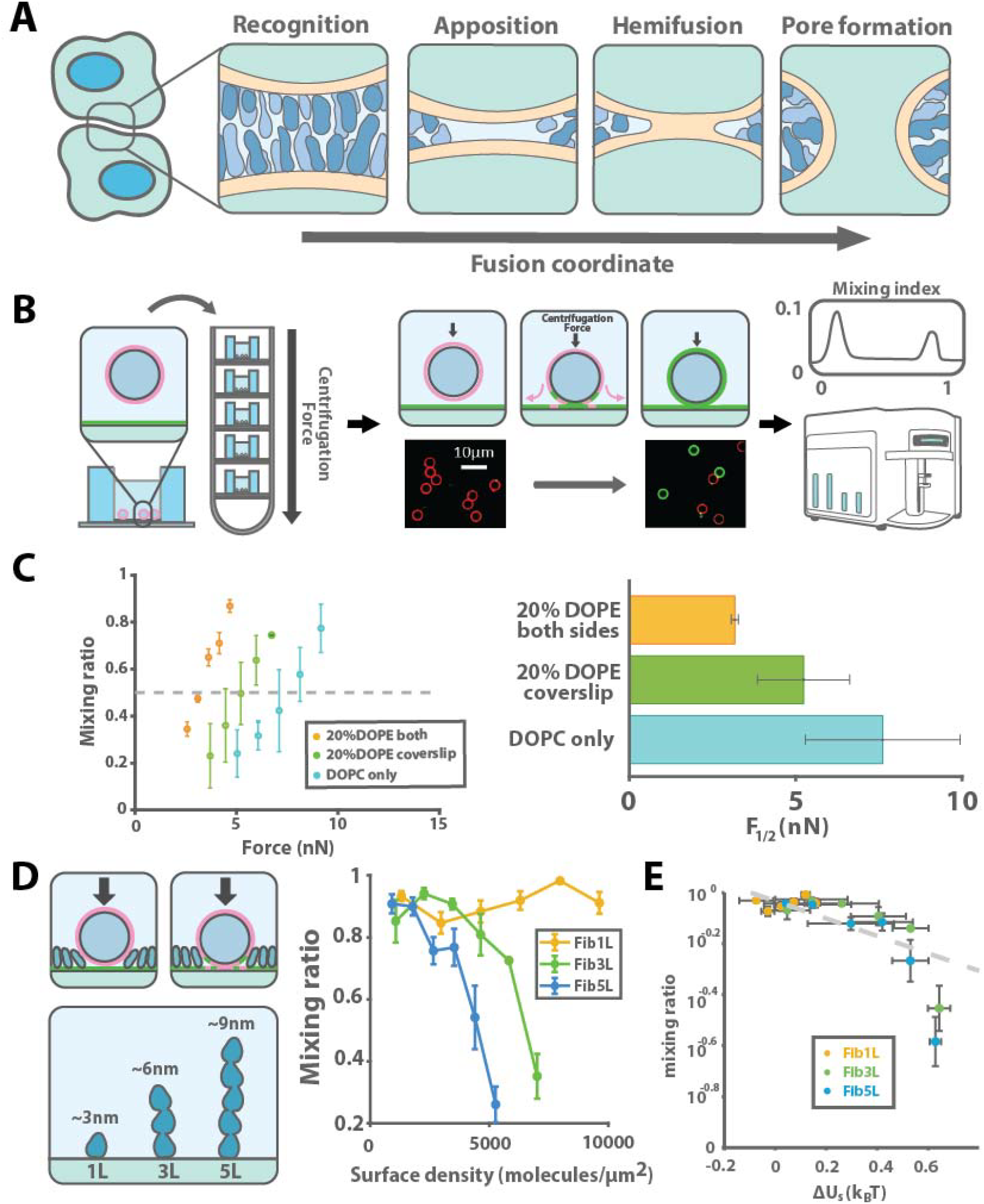
Surface crowding poses a barrier to membrane fusion *in vitro*. **A**, Cell-cell fusion requires close membrane apposition and surface protein displacement. **B**, Our reconstituted membrane fusion assay applies force via centrifugation to drive membranes (red) on a bead in solution and on a surface (green) together. The mixing ratio, defined as the ratio of the surface to bead SLB fluorescence (Liss Rhodamine PE and Atto488PE respectively) between the bead and surface bilayers is dependent on the force applied and lipid composition of the respective surfaces. Linear interpolation gave an estimate of *F*_l/2_, which was compared between three lipid conditions. Error bars represent standard deviation (SD) in mixing ratio over 2-3 replicates and were propagated by linear interpolation from experimental error. **C**, To test the effect of surface protein composition on membrane fusion, the planar SLB was decorated with purified proteins of varying size and fused to beads with a constant force of 5nN. **D**, The mixing ratio was calculated for surfaces decorated with varying densities of Fibcon repeats containing 1, 3, or 5 domains (Fib1L - yellow, Fib3L - green and Fib5L - blue lines). Error bars reflect SD over 2-3 technical replicates. **E**, Mixing ratio for each protein density and type was compared to the crowding energy for the corresponding condition to collapse the data, which was fit to a linear model. Horizontal error bars reflect standard error of the mean (SEM) over 3 replicates for crowding measurements, while vertical ones reflect SD of mixing ratio.

The eukaryotic cell surface is decorated with a forest of glycosylated proteins and lipids, known as the glycocalyx^15,16^ at densities on the order of 20,000 molecules per square micron^17^. Because these surface glycoproteins range in height from <1nm to nearly 1000nm tall^18^, surface density alone is a poor metric of the steric barrier that could resist local exclusion of surface proteins, hindering cell-cell fusion. We recently developed a method to quantify the steric barrier presented by the cell surface, which we refer to as crowding, by measuring the binding affinity (or dissociation constant) of an antibody for an exogenous sensor molecule integrated into a lipid membrane^19^. We showed that the dissociation constant of an antibody for the sensor on a live cell surface is typically 10 times or more greater than it is for the crowding sensor on a bare surface, corresponding to a 2− 3*k*_*B*_*T* barrier against antibody binding created by cell surface molecules^19^. This suggests that cell surface crowding could be a nontrivial but underappreciated barrier to cell-cell fusion, one that could potentially be modulated to regulate fusion rates.

In this work, we combine cell surface crowding measurements with in vitro reconstitution and live-cell fusion assays to demonstrate that cell surface crowding significantly reduces the efficiency of cell-cell fusion. We first develop an in vitro fusion assay to measure the force required to fuse reconstituted membranes and show that adding surface proteins slows fusion in a manner quantitatively consistent with the steric barriers they impose. We then demonstrate that this principle applies in PEG-mediated fusion assays, where cells with less crowding showed greater likelihood of fusion. In the case of cells expressing viral fusogens, we find that differences in cell surface crowding can result in the selective fusion of less crowded cells, spatially segregating crowded (unfused) cells away from multinucleated syncytia of less crowded cells. Finally, we demonstrate that surface crowding is naturally reduced in differentiating myoblasts as they approach a fusogenic state, suggesting that surface crowding can be modulated by cells in physiological contexts to control the timing of cell-cell fusion.

## Results

### A reconstituted membrane mixing assay demonstrates the dependence of fusion on membrane surface crowding

We first sought to investigate the relationship between surface crowding and membrane fusion by developing an *in vitro* assay to quantify lipid mixing, a step toward full membrane fusion, as a function of force. To collect large numbers of events simultaneously, we used centrifugation to exert a compressive force between 3.8-micron glass beads coated with supported lipid bilayers (SLBs) and an SLB-coated glass coverslip (Fig. 1B). The SLBs were composed of DOPC (1,2-dioleoyl-sn-glycero-3-phosphocholine), with the bead SLB labeled with Liss-Rhodamine PE and the coverslip SLB labeled with Atto488-PE, such that lipid mixing between the membranes due to force resulted in the simultaneous loss of Liss-Rho and gain of Atto 488 signal around the bead after 20 minutes of centrifugation. The ratio of Liss-Rho to Atto fluorescence was quantified by flow cytometry, giving a generally bimodal population (Fig. S1), which was used to calculate the mixing ratio, i.e., the fraction of beads found to have undergone lipid mixing for a given force. We then calculated the *F*_l/2_, the force required to give a mixing ratio of 0.5, where membrane fusion had occurred in 50% of the beads in the assay (Fig. 1C). To validate that this approach captures known effects of specific lipid species on membrane fusion, we compared the DOPC membranes to SLBs containing 20% DOPE (1,2-Dioleoyl-sn-glycero-3-phosphoethanolamine), a lipid that induces positive curvature. We found that including DOPE in the coverslip membrane decreased *F*_l/2_ from 7.6 ± 2.3nN to 5.2 ± 1.3nN, while including DOPE in both the bead and coverslip membranes further decreased *F*_l/2_ to 3.17 ± 0.1nN, consistent with previous work showing that DOPE increases lipid mixing^20^ (Fig. 1C).

Next, to quantify the effect of surface crowding on membrane fusion, we decorated the coverslip SLB with purified His-tagged proteins formed from repeats of fibronectin type III consensus (Fibcon) domains using Ni-NTA lipids (Fig. 1D), which we had previously used to create synthetic surface proteins with variable heights^21^. We found that the single Fibcon domain protein (Fib1L) was insufficient to significantly change the mixing ratio, up to the maximum surface density allowable by the SLB without disrupting the lipid bilayer. In contrast, decoration with the triple Fibcon domain protein (Fib3L) decreased the mixing ratio 50% at a density of around 6500 molecules/µm^2^, and only about 4500 molecules/µm^2^ of the quintuple Fibcon (Fib5L) domain protein was required to reduce the mixing ratio to 50% (Fig. 1D). These results suggest that not only the density but also the size of the surface protein is important for its ability to inhibit fusion.

We hypothesized that the steric barrier presented by cell surface crowding could directly explain the decrease in membrane fusion *in vitro*. We previously showed that the energy penalty for insertion of a molecule into a polymer brush, here defined as *ΔU*_s_, can be read out by measuring the apparent dissociation constant (*K*_d_) of an adsorbed molecule for the surface in question compared to a bare surface (*K*_d_^0^)^19^, where *K*_d_ =*K*_d_ ^0^ exp (*ΔU* /*k*_B_*T*). Since the fusion interface must be entirely cleared of both bystander and solvent molecules for fusion to proceed, the fusion rate should simply be proportional to the crowding energy *ΔU*_s_, if crowding is the limiting step to fusion. Using previous measurements of *K*_d_/*K*_d_^0^ for Fibcon domain repeat proteins on SLB-coated beads (Fig. S1), we estimated *ΔU*_s_ and compared it to the mixing ratio for each surface density and protein size condition. This revealed a strong correlation between mixing ratio and crowding energy (Fig. 1E), suggesting that surface crowding quantitatively alters membrane mixing in our reconstituted assay.

### Lowering surface crowding enhances PEG-mediated fusion

Next, we investigated the effect of cell surface crowding on fusion of live cells. We used polyethylene glycol (PEG) to drive cell-cell fusion in the absence of any fusogen, a common practice for generating hybridomas for monoclonal antibody production (Fig. 2A)^22–24^. Adding a high concentration of soluble PEG to the media results in an increase in osmotic pressure between cells, bringing them together and driving fusion. To test whether soluble PEG could drive fusion between cells and SLBs, we dissociated HEK239T cells using 2mM EDTA in PBS and stained them with CellMask Far Red. After allowing them to settle on an Atto488PE-containing SLB coated coverslip, we added soluble PEG to the settled cells and analyzed lipid mixing by flow cytometry after 20 minutes as previously described. We found that 20% PEG8K, was sufficient to cause lipid mixing between the HEKs and surface SLB (Fig. 2C). Gently trypsinizing the HEKs to lower surface crowding (Fig. 2B) prior to incubation with the PEG-containing solution further increased the degree of mixing observed, suggesting that digesting surface proteins drove lipid mixing (Fig. 2C).

**Fig. 2.**
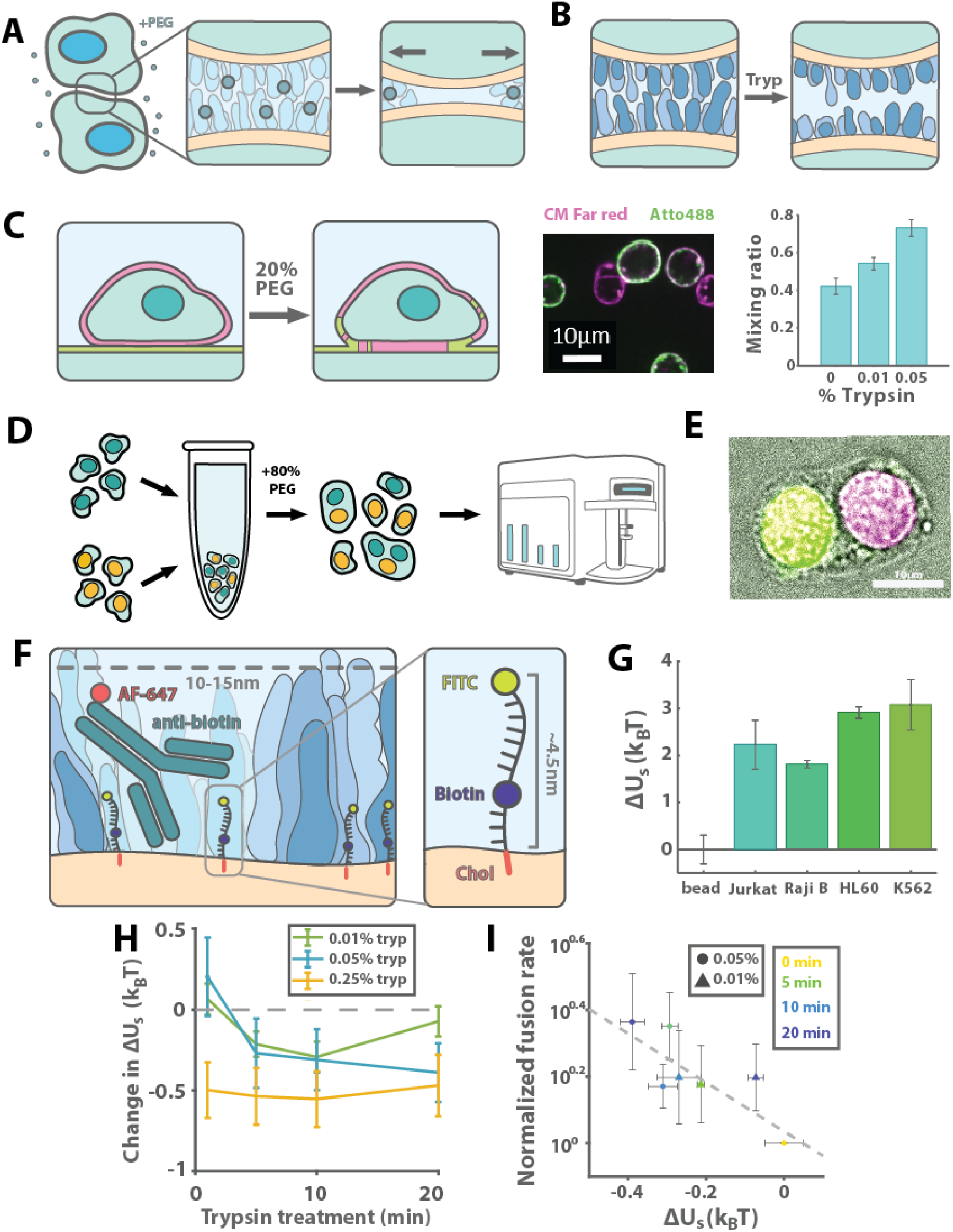
Protein crowding is a barrier to PEG-mediated cell-cell fusion. **A**, PEG was used to exert an osmotic force to fuse cells. **B**, Trypsinization can be used to nonspecifically lower surface crowding. **C**, Centrifugation in the presence of PEG was used to fuse HEK cells (labeled with CM Far Red) to planar SLBs (labeled with Atto488). Cells dissociated with trypsin showed more lipid mixing. Error bars reflect SD over 3 biologically independent replicates. **D**, Cells were fused by incubating suspension cell lines in 80% PEG solution and analyzed by microscopy or flow cytometry. **E**, Jurkat cells expressing H2B-miRFP670 and H2B-GFP were used to validate this approach. **F**, Cell surface crowding can be quantified by incorporating a probe consisting of a DNA oligo decorated with cholesterol, biotin, and fluorescein into cell membranes and measuring binding of an anti-biotin antibody. **G**, *ΔU*_s_ was compared between an SLB-coated bead and four common suspension cell lines. Error bars reflect fit SEM. **H**, Gentle trypsinization of K562 cells was used to vary surface crowding energy, which was measured with the crowding sensor in flow cytometry and normalized to nontreated control (dashed grey line). Error bars reflect 95% confidence interval for each condition. **I**, Fusion rates compared to the crowding energy changes due to a range of trypsinization conditions in K562 hybridomas and fit to a linear model (dashed line; *p*<0.05 by F test against a constant model). Horizontal error bars reflect fit SEM for surface energy calculation; vertical error bars reflect SEM for 3-5 biologically and technically independent replicates of PEG-mediated fusion assay.

To test the effect of cell surface crowding on content mixing, which occurs with full fusion rather than only lipid mixing, we developed a PEG-mediated fusion assay for suspension cells. Briefly, we pelleted cells with soluble PEG and quantified the number of fused cells using an imaging flow cytometer and an automated segmentation pipeline (Fig. 2D). Using Jurkat cells, half expressing H2B-GFP and the other half expressing H2B-miRFP, we found that PEG-mediated fusion and content mixing was successful with soluble PEG3.5K (Fig. 2E).

We next tested whether changes in cell surface crowding would alter cell-cell fusion in the PEG-mediated fusion assay. We chose to test K562 cells, which we previously identified as naturally highly crowded, with an energy barrier of about 3*k*_*B*_*T* (Fig. S3), using our crowding sensor (Fig. 2F,G). To reduce crowding by cleaving surface proteins, we incubated cells in dilute trypsin, subsequently quenching with FBS-containing growth media. We found that the decrease in *ΔU*_s_ saturated immediately with 0.25% trypsin, while it reached a similar level with 0.05% trypsin after 20 minutes of treatment, and did not saturate over the same timeframe for the 0.01% trypsin condition (Fig. 2H). We then tested the effect of these treatments on cell-cell fusion with the 0.01% and 0.05% trypsin conditions, reasoning that these conditions would provide the largest range of crowding levels. Combining the fusion efficiency and surface energy data, we find that the log of the fusion probability decayed with the crowding as expected for changes in surface accessibility alone (Fig. 2I; Supplementary Information), suggesting that crowding may be a limiting factor for cell-cell fusion in some contexts.

### FAST-protein mediated fusion is inhibited by surface crowding

Having determined that cell surface crowding is a barrier to PEG-mediated fusion, we wondered whether it is also a barrier to fusogen-mediated cell-cell fusion. We utilized the fusion associated small transmembrane (FAST) protein p14, which, despite its 1.5nm height^25,26^, is able to drive fusion by recruiting the actin cytoskeleton^27,28^. We previously showed that overexpression of tall proteins, such as the Signal-Regulatory Protein Alpha (SIRPα) ectodomain (∼14nm tall^29^), can increase surface crowding, while overexpression of small proteins, such as Fib1L (∼3nm tall), has little to no effect^19^ (Fig 3A). Thus, to test the effect of crowding on p14-mediated fusion, we co-transfected either the tall SIRPα or the short Fib1L, which act as ‘bystanders’ since they are not required for fusion, with p14 and quantified cell-cell fusion by fluorescence microscopy (Fig 3B). Strikingly, we found that the tall bystander nearly completely inhibited fusion, while the short bystander had little effect compared to an mCherry transfection control (Fig. 3C).

**Fig. 3.**
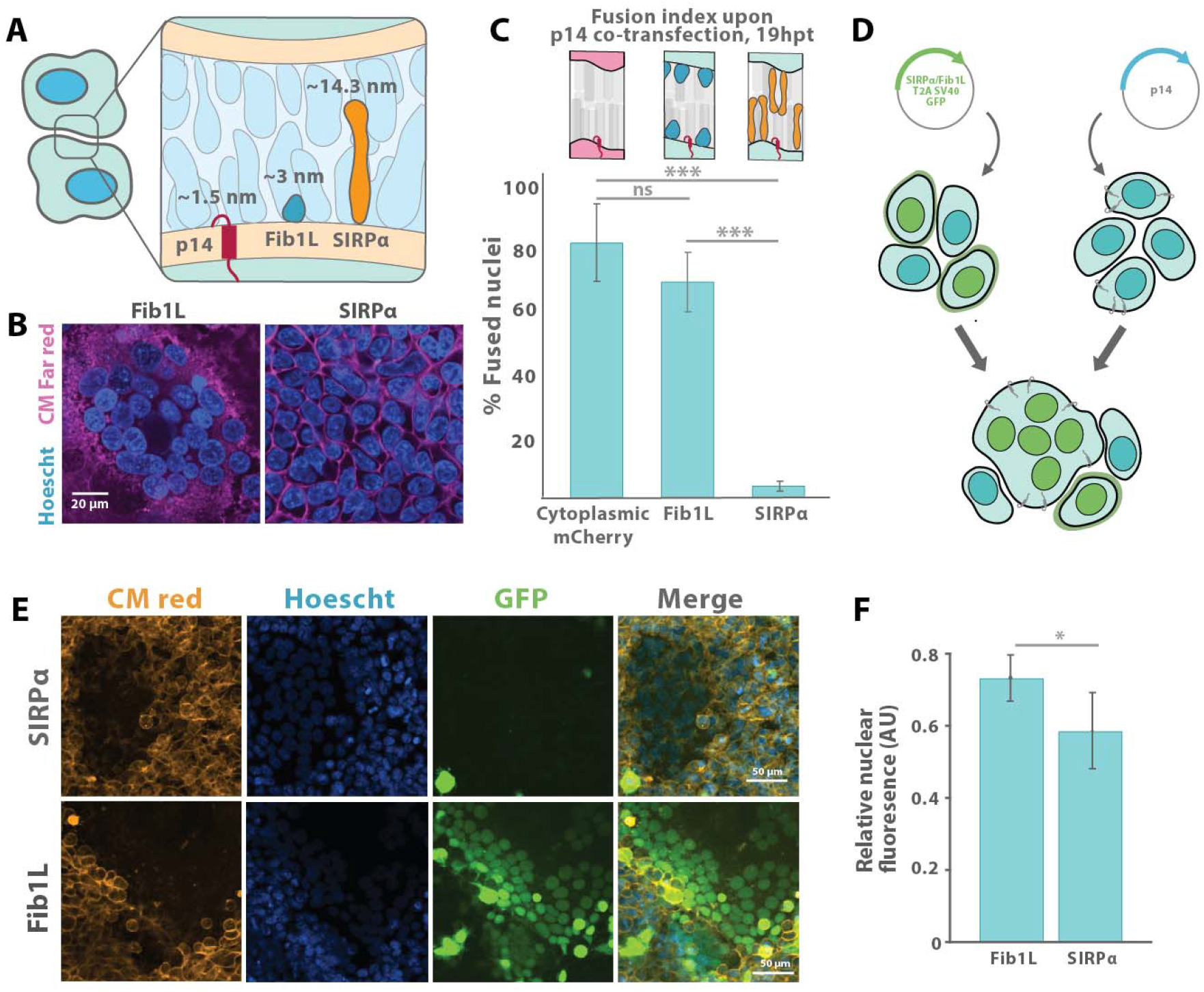
Increased surface crowding inhibits p14-mediated cell-cell fusion, enabling fusion selectivity. **A**, The FAST protein p14 has been estimated to be 1.5nm in height, considerably shorter than the bystanders used here. **B**, HEK293 cells were co-transfected with p14 and Fibcon1L, SIRP□ ectodomain, or cytoplasmic mCherry control. **C**, For each condition, the fraction of all nuclei found in syncytia was calculated and averaged over three biologically and technically independent replicates. Fib1L and SIRPα were both significantly less fusogenic than mCherry control (*p*<0.01 by two-tailed T test with Bonferroni correction). Error bars reflect SEM. **D**, The rate of fusion of individual transfected cells was assessed by designing nuclear reporter constructs consisting of an ectodomain and a polycistronic fluorescent protein with an NLS. **E**, The NLS-GFP spread between all nuclei within a particular syncytium following fusion for both constructs. **F**, The mean nuclear fluorescence was calculated within syncytia, divided by the average nuclear fluorescence per image to normalize for transfection efficiency, and compared between constructs. The propensity of cells expressing the SIRPα ectodomain to be fused into syncytia was less than that of cells expressing Fib1L (*p*<0.05 by two-tailed T test). Error bars reflect SEM over three biologically independent replicates.

Based on this result, we wondered whether cell-cell fusion within a population of cells would be biased towards less crowded cells, potentially conferring a degree of specificity and spatial control over the fusion process based only on surface energy barriers. To test this, we designed a synthetic bystander protein with a nuclear reporter, consisting of a SPOT-tagged bystander ectodomain (i.e., either Fib1L or SIRPα) with a downstream T2A cleavage site and an eGFP-tagged SV40 nuclear localization signal (NLS) (Fig. 3D). We observed the expected surface localization of the SPOT-label and nuclear localization of the GFP signal by microscopy, and we confirmed proportionality between GFP expression level and SPOT-label binding by flow cytometry (Fig. S3).

We quantified the bias in fusion by co-culturing HEK cells expressing either Fib1L or SIRPα bystander constructs with HEKs transfected with p14. We observed that once fused, the NLS-GFP signal expressed by a single nucleus would quickly distribute itself across the syncytium (Fig. 3E). We then segmented the cells (Fig. S4) and quantified the fraction of cells expressing the respective ectodomains included in the syncytia by calculating the average GFP fluorescence of nuclei inside the syncytia normalized by the average GFP level across all nuclei. We found this relative fluorescence was approximately 20% lower for cells expressing the SIRPα ectodomain compared to those expressing the Fib1L ectodomain, suggesting that expression of the taller ectodomain indeed biased p14 fusion toward otherwise identical cells with lower energy barriers, an indication that cell surface crowding can guide fusion selectivity (Fig. 3F).

### Cell surface crowding is modulated during myoblast development and fusion

Finally, we asked whether cellular systems might already tune cell surface crowding to regulate fusion in a physiological context. We considered the process of muscle development, a highly regulated and essential process in vertebrates wherein progenitor cells differentiate and fuse into myotubes (Fig. 4A). C2C12 murine myoblasts, a well-characterized cell culture model for muscle development^30^, elongate after serum starvation and fuse into large syncytial fibers after 4-6 days in culture (Fig. 4B). We monitored the changes in cell surface crowding with our crowding sensor throughout this process of differentiation, dissociating the myoblasts with EDTA and quantifying crowding by flow cytometry. Interestingly, we observed a gradual but steady decrease in the level of surface crowding throughout the differentiation process, with a change in the surface crowding energy barrier of 1*k*_*B*_*T* between 0 and 6 days of differentiation (Fig. 4C), indicating that the cell surface becomes significantly less crowded as differentiation proceeds.

**Fig. 4.**
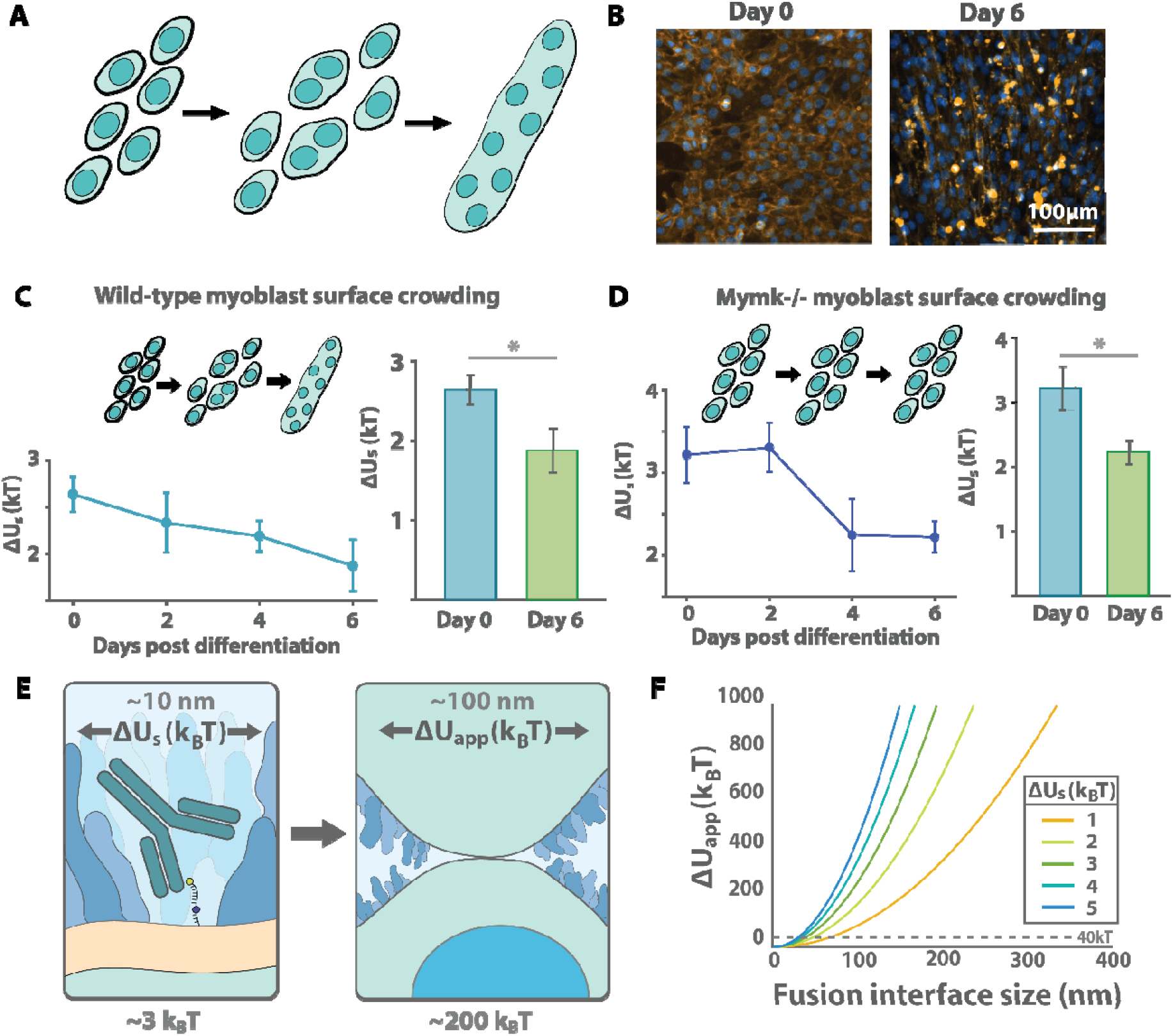
Myoblasts reduce their cell surface crowding as they differentiate and undergo fusion. **A**, The C2C12 line is a model for myoblast differentiation, during which cells elongate and then fuse. **B**, Following serum starvation, wild type C2C12 cells were stained with CellMask Orange and Hoechst 33342, demonstrating myotube formation on day 6. **C**, Following serum starvation, crowding was measured in wild C2C12 cells using the aforementioned crowding probe at two-day intervals. Error bars reflect SEM over three biologically independent replicates (*left*). Days 0 and 6 were directly compared and found to be significantly different (p<0.05). **D**, Crowding was measured in Mymk^-/-^ C2C12 cells following serum starvation. **E**, The crowding sensor measures the energy to clear an IgG-sized patch of membrane. We extrapolate this measurement based on parameters from the literature to estimate for membrane apposition for a 100 nm fusion pore. **F**. Full extrapolation of crowding sensor measurement to the energetic cost of membrane apposition, (see Supplementary Information).

To confirm that our measurements were not influenced by syncytia formation itself, which may reduce crowding due to surface-volume ratios and other effects, we also measured crowding in a C2C12 knockout line lacking the myoblast fusogen Myomaker (Mymk^-/-^), which is necessary for cell-cell fusion^31^. Consistent with previous work, we observed elongation and morphological changes consistent with differentiation upon serum starvation, but no fusion (Fig. S4). Repeating our crowding sensor measurement on the knockout line revealed a similarly significant decrease in crowding between days 0 and 6 (Fig. 4D). This suggests that the observed crowding changes are independent of syncytia formation and are instead modulated by the differentiation process itself.

## Discussion

### Surface crowding presents a potentially rate-limiting barrier to fusion

In this work, we quantitatively demonstrate that surface crowding hinders cell-cell fusion by presenting a physical barrier to close membrane apposition. We observed that lipid mixing ratio and cell-cell fusion rate were both strongly correlated with surface crowding energy, suggesting that surface crowding dominates the energetics of fusion both *in vitro* and *in vivo*. This data suggests that in physiological membrane fusion, exclusion of surface proteins presents a potentially rate-limiting step in the fusion pathway.

By directly quantifying cell surface crowding, we can estimate the energetic penalty of excluding cell surface proteins during cell-cell fusion. From our measurements showing a penalty of *ΔU*_s_ 2 ≈ 3*k*_*B*_*T* for incorporation of an IgG antibody, with dimensions of ∼10-15nm^32^ (Fig. 4E), we estimate the energetic cost of clearing a 100nm diameter fusion interface to be ∼200*k*_*B*_*T* (Supplementary Information; Fig. 4F), exceeding the 40*k*_*B*_*T* barrier estimated for bare membrane fusion. The presence of a robust cell surface barrier preventing neighboring cells from fusing is rather reassuring; spurious fusion of cells in close contact could lead to apoptosis or other anomalous behavior that must be avoided in multicellular organisms. By controlling the considerable energetic barriers presented by surface crowding, cells can dramatically lower the possibility of cell-cell fusion.

### Modulation of the cell surface may enable control of fusion dynamics

Considering the large barrier cell surface crowding presents to fusion, small changes in crowding could result in large changes in rates of cell-cell fusion. This idea is empirically supported by the observation that gentle trypsinization results in a 40% increase in PEG-mediated fusion, and the 1*k*_*B*_*T* increase in crowding energy due to SIRPα Overexpression^19^ nearly abrogated p14-driven fusion. Similarly, we showed that the cell surface of developing myoblasts is reduced by 1*k*_*B*_*T* following differentiation. In either of these instances, the energetic barrier to fusion is decreased by as much as 50%. Taken together, we speculate that the large energetic barrier posed by cell surface crowding, which can be modified enzymatically or by turnover of the cell membrane, makes this biophysical barrier a biologically convenient mechanism by which cells modulate fusion.

In summary, we have shown through *in vitro and in vivo* examples, with and without fusogens, that cell surface crowding is an important constraint on the process of cell-cell fusion, one that could be harnessed for cell engineering and delivery.

## Acknowledgements

We would like to thank Pengpeng Bi (University of Georgia) for the myomaker knockout cell line and useful discussions. We would also like to thank Mary West of the QB3 Cell and Tissue Analysis Facility for flow cytometry assistance, and the members of the Fletcher lab for useful discussions. D.A.F. is supported by NIH R01 GM134137, the NSF Center for Cellular Construction (DBI-1548297), and the Miller Institute for Basic Research. D.A.F. is a Chan Zuckerberg Biohub Investigator.

## Author Contributions

D.S.W.L., L.F.O., S.S., and D.A.F. designed the research; D.S.W.L., L.F.O., and S.S. performed experiments and formal analysis. D.S.W.L., L.F.O., and D.A.F wrote the paper.

## Data and Code Availability

All data and code associated with this manuscript are available at the Center for Open Science, at DOI 10.17605/OSF.IO/AF2Q7, and any further requests may be directed to the corresponding author.

## Reagent Availability

All reagents associated with this manuscript are available upon request to the corresponding author.

## Competing interests

The authors declare no competing interest.

## Notes

https://osf.io/af2q7/

### Competing Interest Statement

The authors have declared no competing interest.

