## Supplementary material for "Cell surface crowding is a tunable biophysical barrier to cell-cell fusion": Methods and Supplementary Information

#### Bead preparation and centrifugation

*Bead and coverslip cleaning*

5um silica microspheres (Bangs Laboratories, Inc, SS05003) were washed in a glass test tube with piranha mixture by adding a 2:3 ratio of 30% hydrogen peroxide to full-strength sulfuric acid. The bead and piranha mixture was then sonicated in a bath sonicator for 30 minutes and washed by centrifugation (1000*g*) and resuspended in ultrapure MilliQ water (EMD Millipore). 18x18-1.5mm glass coverslips were mounted onto a teflon rack and RCA cleaned by sequentially boiling them at 80C with a mixture of 5g KOH, 80mL MilliQ water, and 20mL of 30% hydrogen peroxide for 30 minutes in a glass beaker. The teflon rack was then briefly rinsed in a beaker of milliQ water and transferred to another glass beaker of 25mL full-strength HCl, 80mL milliQ water, and 20mL 30% hydrogen peroxide and again boiled at 80°C for 30 minutes. Following the acid wash, the teflon rack was then transferred sequentially into two beakers each filled with milliQ water, and stored in a third sealed beaker of milliQ water for later use.

*Supported lipid bilayer formation*

To prepare the lipids, small unilamellar vesicles (SUVs) were formed by mixing the desired lipid composition (POPC, DOPE from Avanti Polar Lipids) in molar ratios in a round-bottom glass tube (0.28 µmol lipid total), and then simultaneously vortexing and desiccating with nitrogen glass to form a thin film at the bottom of the glass tube. To bind the surface proteins, NiNTA-DGS lipid (Avanti) was included in this initial mixture at a molar concentration of 0.1%-7% . SUVs intended for supported lipid bilayer (SLB) formation included 0.01% Atto-488PE, and SUVs intended to coat the glass beads included 0.01% Liss-rhodamine dye. The glass tubes were then incubated in a vacuum-sealed chamber overnight to ensure evaporation of the chloroform. The dried lipid film was then reconstituted in 900µL of deionized water, then sonicated with a tip sonicator for 3 minutes on ice with a 2 second duty cycle, amplitude 20%. These SUVs were then stored at 4°C in 1x MOPS buffer (adding 100µL of 10X MOPS to the sonicated solution).

To form the bilayers, the SUVs were incubated with the washed glass, allowing them to crash out onto the glass and form a bilayer. Beads were prepared as previously described[1]. Briefly, 40µL of beads were incubated with 20µL of silica microspheres in 160µL MOPS buffer on a rotisserie at room temperature for 20 minutes. Afterwards, the beads were washed 3x via centrifugation at 50*g* and resuspended in MOPs buffer. SLBs were prepared as described previously[2]. Briefly, SUVs were incubated for 15 minutes in a PDMS well situated on a cleaned glass slide. The SLBs were then washed 3x with HEPES buffer. To add proteins to the surface,

SLBs containing NiNTA-DGS lipids were incubated with purified His-tagged Fibcon extracellular domain for 15 minutes at room temperature, then washed 3x with MOPS buffer.

*Lipid mixing by centrifugation*

Custom centrifuge inserts were engineered and 3D printed to hold the glass coverslips horizontally in a Type 70 Ti Fixed-Angle centrifuge rotor. 10µL of lipid-coated beads were added to each SLB well, and the glass slides were mounted on the custom centrifuge inserts. The centrifuge was then spun at a constant speed for 20 minutes. The beads were vigorously pipetted to resuspend all beads, including ones that were fused to the SLB, and the subsequent supernatant was run on the Attune CytPix flow cytometer. Mixing ratio was then calculated by the fraction of beads that exhibited a gain of green signal, indicating that the green SLB lipids mixed with the lipids coating the bead surface.

**Cell culture and maintenance**

All cell lines were acquired from the Barker Hall Cell Culture Facility at UC Berkeley except for C2C12 knockouts, which were a kind gift from Pengpeng Bi (University of Georgia). HEK and C2C12 cells were cultured in T25 cell-culture treated flasks in DMEM media supplemented with GlutaMax, 10% FBS, and 1% pen-strep. Jurkat and HL60 cells were cultured in RPMI with 10% FBS and 1% pen-strep. K562 and Raji B cells were cultured in RPMI with 10% FBS and 1% pen-strep and 1% sodium pyruvate. Cells were split every two days at 1:5 and kept in an incubator at 37°C and 5% $CO_2$.

C2C12 cells were differentiated by removing growth media, washing with PBS, and replacing differentiation media (DMEM containing 2% horse serum and 1% pen-strep). Cells were gently washed, and media was replaced every 24 hours following initial differentiation.

**Plasmid generation**

For all plasmids, gene blocks were ordered from Twist Biosciences and  inserted by Gibson assembly (New England Biolabs) into linearized and gel purified (Qiagen)  pHR vectors either containing only the plasmid backbone or previously used ectodomains.[3] Plasmid sequences were confirmed by Sanger sequencing over the insert or full plasmid Nanopore sequencing (UC Berkeley Sequencing Facility at Barker Hall)..

Gene block sequences are as follows:

| | |
|---|---|
| T2A-SV40NLS-GFP | CGGATCAAACAGAAGAAAGCCAAGGGGT<br>CAGAGGGCAGAGGAAGTCTGCTAACATG<br>CGGTGACGTCGAGGAGAATCCTGGCCCA<br>CCAAAGAAGAAGCGTAAGGTAATGGTGA<br>GCAAGGGCGAGGAGCTGTTCACCGGGGT<br>GGTGCCCATCCTGGTCGAGCTGGACGGC<br>GACGTAAACGGCCACAAGTTCAGCGTGT |

| | |
|---|---|
| | CCGGCGAGGGCGAGGGCGATGCCACCT<br>ACGGCAAGCTGACCCTGAAGTTCATCTGC<br>ACCACCGGCAAGCTGCCCGTGCCCTGGC<br>CCACCCTCGTGACCACCCTGACCTACGG<br>CGTGCAGTGCTTCAGCCGCTACCCCGAC<br>CACATGAAGCAGCACGACTTCTTCAAGTC<br>CGCCATGCCCGAAGGCTACGTCCAGGAG<br>CGCACCATCTTCTTCAAGGACGACGGCAA<br>CTACAAGACCCGCGCCGAGGTGAAGTTC<br>GAGGGCGACACCCTGGTGAACCGCATCG<br>AGCTGAAGGGCATCGACTTCAAGGAGGA<br>CGGCAACATCCTGGGGCACAAGCTGGAG<br>TACAACTACAACAGCCACAACGTCTATAT<br>CATGGCCGACAAGCAGAAGAACGGCATC<br>AAGGTGAACTTCAAGATCCGCCACAACAT<br>CGAGGACGGCAGCGTGCAGCTCGCCGAC<br>CACTACCAGCAGAACACCCCCATCGGCG<br>ACGGCCCCGTGCTGCTGCCCGACAACCA<br>CTACCTGAGCACCCAGTCCGCCCTGAGC<br>AAAGACCCCAACGAGAAGCGCGATCACA<br>TGGTCCTGCTGGAGTTCGTGACCGCCGC<br>CGGGATCACTCTCGGCATGGACGAGCTG<br>TACAAGTAAGCGGCCGCGACTCTAGAGT<br>CGACCTGCAGGCATGCAAGCTTGATATCA<br>AGCTTATCGA |
| T2A-SV40NLS-miRFP | CGGATCAAACAGAAGAAAGCCAAGGGGT<br>CAGAGGGCAGAGGAAGTCTGCTAACATG<br>CGGTGACGTCGAGGAGAATCCTGGCCCA<br>CCAAAGAAGAAGCGTAAGGTAATGGTAGC<br>AGGTCATGCCTCTGGCAGCCCCGCATTC<br>GGGACCGCCTCTCATTCGAATTGCGAACA<br>TGAAGAGATCCACCTCGCCGGCTCGATC<br>CAGCCGCATGGCGCGCTTCTGGTCGTCA<br>GCGAACATGATCATCGCGTCATCCAGGC<br>CAGCGCCAACGCCGCGGAATTTCTGAAT<br>CTCGGAAGCGTACTCGGCGTTCCGCTCG<br>CCGAGATCGACGGCGATCTGTTGATCAA<br>GATCCTGCCGCATCTCGATCCCACCGCC<br>GAAGGCATGCCGGTCGCGGTGCGCTGCC<br>GGATCGGCAATCCCTCTACGGAGTACTG<br>CGGTCTGATGCATCGGCCTCCGGAAGGC<br>GGGCTGATCATCGAACTCGAACGTGCCG<br>GCCCGTCGATCGATCTGTCAGGCACGCT<br>GGCGCCGGCGCTGGAGCGGATCCGCAC<br>GGCGGGTTCACTGCGCGCGCTGTGCGAT<br>GACACCGTGCTGCTGTTTCAGCAGTGCAC<br>CGGCTACGACCGGGTGATGGTGTATCGT<br>TTCGATGAGCAAGGCCACGGCCTGGTATT<br>CTCCGAGTGCCATGTGCCTGGGCTCGAA |

TCCTATTTCGGCAACCGCTATCCGTCGTC
GACTGTCCCGCAGATGGCGCGGCAGCTG
TACGTGCGGCAGCGCGTCCGCGTGCTGG
TCGACGTCACCTATCAGCCGGTGCCGCT
GGAGCCGCGGCTGTCGCCGCTGACCGG
GCGCGATCTCGACATGTCGGGCTGCTTC
CTGCGCTCGATGTCGCCGTGCCATCTGC
AGTTCCTGAAGGACATGGGCGTGCGCGC
CACCCTGGCGGTGTCGCTGGTGGTCGGC
GGCAAGCTGTGGGGCCTGGTTGTCTGTC
ACCATTATCTGCCGCGCTTCATCCGTTTC
GAGCTGCGGGCGATCTGCAAACGGCTCG
CCGAAAGGATCGCGACGCGGATCACCGC
GCTTGAGAGCTAAGCGGCCGCGACTCTA
GAGTCGACCTGCAGGCATGCAAGCTTGA
TATCAAGCTTATCGA

**Lentiviral preparation and cell line generation**

Lentivirus was produced by transfecting HEK293T cells with the plasmid of interest, pCMV-dR8.91, and pMD2.G (1.5µg, 1.33µg, and 0.167µg per 35mm well) using Mirus TransIT-293 Transfection Reagent per manufacturer's protocol. After 60-72 hours, supernatant containing viral particles was harvested and filtered with a 0.45 µm syringe filter (Corning). Supernatant was immediately used for transduction or aliquoted and stored at −80°C. Cells were seeded at 20% confluency in 35mm dishes and 0.1-1mL of filtered viral supernatant was added to the cells. Media containing virus was replaced with fresh growth medium 24 hr post-infection. Infected cells were imaged to assess transduction efficiency and then used in flow cytometry assays as described above.

**PEG-mediated cell-cell fusion assay**

Cells were counted and mixed at 500,000 cells per condition. Each mixture was then centrifuged and washed 2x in serum-free media at 500*g* for 5 minutes. Cells were then pelleted, and 500µL of PEG3.5k (80% w/v in PBS) was added to the pellet and incubated at room temperature for 2 minutes. 10mL of serum-free media was then added to the mixture, and cells were allowed to recover at room temperature for 30 minutes. The pellet was then washed 2-3 times with serum-free media until the pellet became loosened. The pellet was carefully resuspended in 1mL of serum-free media and transferred to a 96-well plate to quantify fluorescence in the Attune CytPix flow cytometer.

Cells were counted and mixed at 500,000 cells per condition. Each mixture was then centrifuged and washed 2x in serum-free media at 500*g* for 5 minutes. Cells were then pelleted, and 500µL of PEG3.5k (80% w/v in PBS) was added to the pellet and incubated at room temperature for 2 minutes. 10mL of serum-free media was then added to the mixture, and cells were allowed to recover at room temperature for 30 minutes. The pellet was then washed repeatedly with serum-free media until the pellet became loosened. The pellet was then carefully resuspended in 1mL of serum-free media and transferred to a 96-well plate to quantify fluorescence in the Attune CytPix flow cytometer.

### p14 cell-cell fusion experiments

*Sample preparation and microscopy*

CellVis 8-well TC-treated chambers were first fibronectin coated by incubating each well with fibronectin diluted 1:50 in PBS for 30m at 37°C. HEK293T cells were plated one day before transfection and seeded at 20,000 cells/well. The following day, they were transfected with Mirus TransIT 293 according to manufacturer protocol. 19 hours post-transfection, cells were incubated with Hoescht and CellMask for 10 minutes at 37°C, and then transferred to the confocal microscope for imaging using a Nikon W1 spinning disk microscope on an Ti2 body through a 20X air objective (E Plan, NA=0.40) with a Orca Fusion BT CMOS camera.

*Image Analysis*

To analyze cell-cell fusion efficiency, at least 1500 nuclei per well were segmented using the CellPose "nuclei" model based on Hoechst labeling. The CellMask channel was used to identify syncytia, either manually for counting experiments or by custom automated MATLAB pipeline for quantifying nuclear expression levels. Each condition was averaged over at least four repeats to determine fusion percentage.

### Crowding sensor measurements

Crowding sensor measurements were performed as previously described[3]. The crowding sensor oligo, 5′-FITC-TTTTTT-biotin-TTT-cholesterol-3′, was ordered from IDT and resuspended at 100$\mu$M. Cells (dissociated with Versene, if necessary) or beads coated with SLBs (assembled as described above and containing a lipid composition of 79% molar fraction DOPC, 1% DOPS, 20% cholesterol from Avanti Polar Lipids) were pelleted and resuspended in PBS. They were then chilled on ice for 15 minutes and incubated with oligo at a concentration of 50nM. Cells were then washed three times by centrifugation and resuspension and incubated with the desired concentration of Alexa Fluor 647-conjugated anti-biotin antibody BK-1/39 (Santa Cruz Biotechnology, sc-53179) at a cell density of ~10000-20000 per 200$\mu$L. Cells were then assayed in a 96-well format using the autosampler of an Attune Cytpix Flow Cytometer with the BRVY configuration and the factory default filters. Data was analyzed using a custom script in MATLAB. The occupancy ratio $\theta$ was calculated by taking the ratio of 647 fluorescence (RL1-H) to 488 fluorescence (BL1-H). The occupancy ratio was then fit to a Langmuir isotherm to calculate the $K_d$, which was normalized to SLB-bead condition.

### Estimation of apposition energy

We developed a scaling argument to describe the relationship between surface crowding and fusion. It has previously been shown that the energy penalty for insertion of a molecule into a polymer brush, here defined as $\Delta U_s$, can be defined as the product of the osmotic pressure of the brush layer, $\Pi(\phi)$, and the effective volume of the inserted molecule, $V_{ab}$, which is an antibody molecule in the case of our sensor. To convert this measured crowding energy to that required to clear a surface of some arbitrary size, e.g., as would be required for fusion, we simply rescale this energy by the relative volume of the fusion site compared to that of the antibody, i.e., $\mathit{\Delta} U_{\mathrm{patch}} = \mathit{\Delta} U_{\mathrm{s}} {L^2}_{\mathrm{fuse}} / {L^2}_{\mathrm{ab}}$. We estimate that the size of the antibody is $L_{\mathrm{ab}} \approx 10 -$ `$5$nm, while we estimate the size of the fusion patch on one membrane to be $L_{\mathrm{fuse}} \approx$ 50-100nm based upon imaging done in the *Drosophila* myoblast[4]. The probability that this occurs at equilibrium will go as the exponential of this energy, according to a standard Boltzmann distribution, i.e., $p = \exp\left(-\mathit{\Delta} U_{\mathrm{patch}} / k_{\mathrm{B}} T\right)$. Assuming that this must occur independently on two apposed surfaces, the probability will scale additively with the crowding energy, and so the energy required for close apposition of two membranes is $\mathit{\Delta} U_{\mathrm{app}} = 2 \mathit{\Delta} U_{\mathrm{s}} {L^2}_{\mathrm{fuse}} / {L^2}_{\mathrm{ab}}$.

### Supplementary Figures

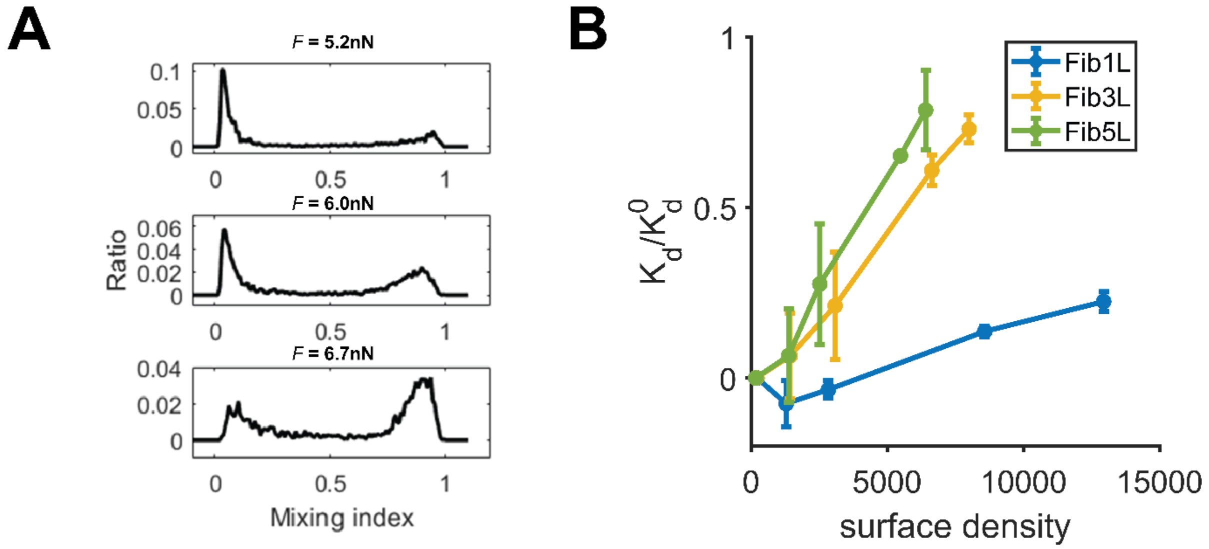


**Fig. S1:** Computation of mixing ratio and crowding interpolation. **A**, The mixing ratio was computed by taking the percentage of beads found in the peak in the histogram corresponding to a higher fluorescence ratio. **B**, The $\Delta U_s$ for Fibcon repeats on beads was computed by using previously published data[3] (blue, yellow, green) and linearly interpolating for the surface densities used in the centrifugation experiments.

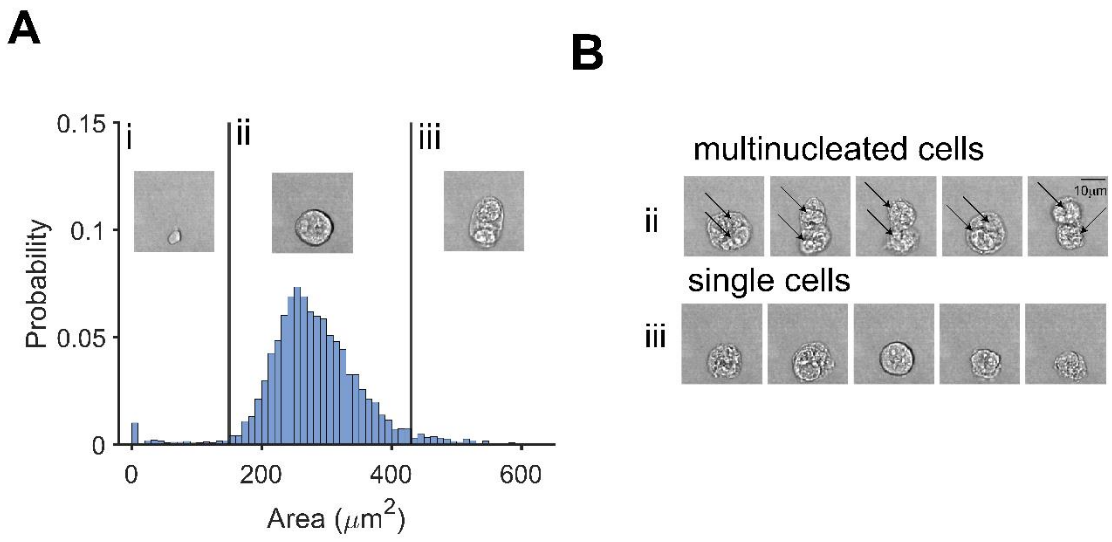


**Fig. S2:** Identifying fused cells by imaging flow cytometry. **A,** High throughput imaging flow cytometry was used to determine the fusion percentage in the hybridoma experiments. Bright field images (*inset*) were taken for each event and segmented, producing a histogram of cell areas. A roughly 2-standard-deviation-threshold was used to separate i) debris, ii) single cells, and iii) multinucleated hybridomas. **B,** This thresholding approach was validated by randomly selecting 5 images in each condition. The events categorized as iii) multinucleated hybridomas all showed multiple nuclei, while the ii) single cells did not but did appear to be intact single cells.

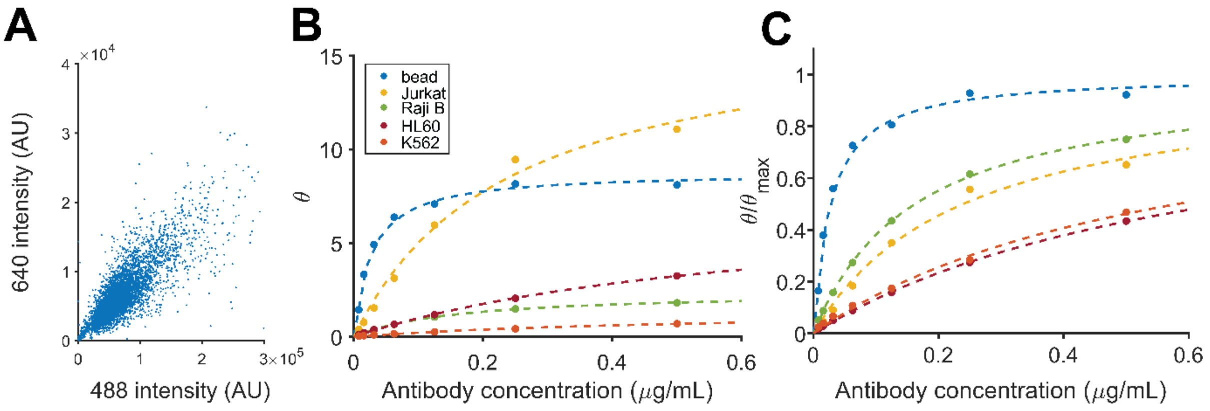


**Fig S3:** Crowding probe analysis. **A,** Example flow cytometry data for the crowding probe. For each antibody/cell condition, the crowding probe occupancy ratio $\theta$ was calculated by taking the ratio of antibody (640 fluorescence) to oligo (488) for each cell. **B,** Occupancy ratio was calculated for a range of antibody concentrations for each cell condition since saturation ratios may differ between cell types. Fluorescent ratios were fit to the Langmuir isotherm, i.e., $\theta = \theta_{\mathrm{max}} \frac{c}{c+K_{\mathrm{d}}}$ (dashed line). **C,** The occupancy ratio, normalized $\theta_{\mathrm{max}}$, i.e., maximum sensor occupancy, across cell types, which we treated as a free parameter, was plotted for each condition for visual comparison of the $K_{\mathrm{d}}$. The $K_{\mathrm{d}}$, was calculated to assess the affinity change due to cell surface crowding and normalized to that of a bead ($K_{\mathrm{d}}^{0}$), giving the calculated crowding energy barrier.

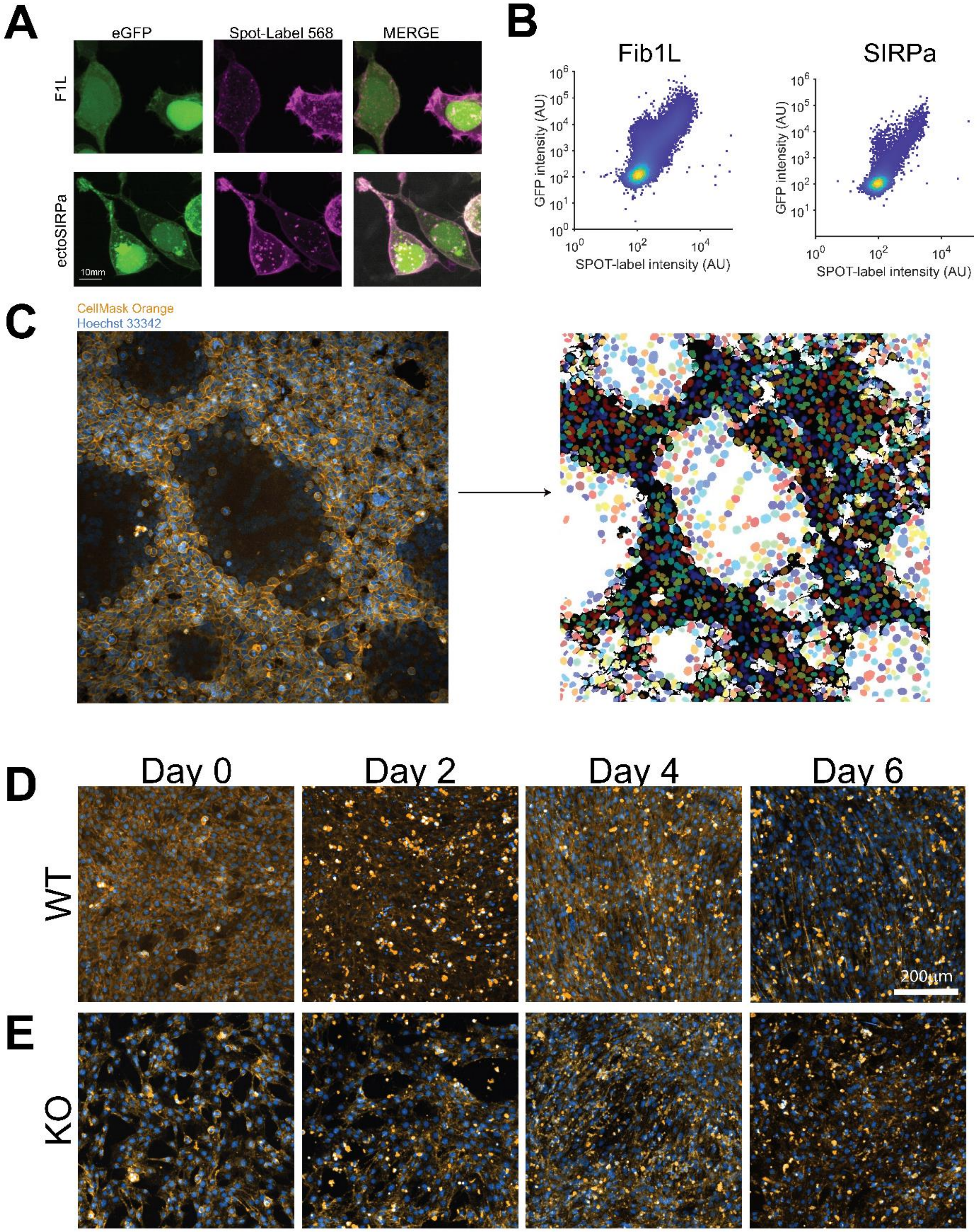


**Fig. S4:** Validation of cell-cell fusion assays. **A,** Transfected cells were stained with SPOT-label 568 and imaged or quantified by flow cytometry. Surface localization of SPOT-label and nuclear localization of fluorescent protein were observed. **B,** A direct proportionality between GFP and spot-label stain intensity were observed by flow cytometry. **C,** For p14 experiments, cells were stained with CellMask Orange and Hoechst 33342 to independently image the cell membrane and nucleus (*left*). Example segmentation of syncytia (black/white) and nuclei (right). Nuclei overlapping the white region were defined as in syncytia and those overlapping the black region were defined as not in syncytia. **D,** Microscopy for validation of myoblast fusion. Following serum starvation, wild type C2C12 cells were stained with CellMask Orange and Hoechst 33342, demonstrating myotube formation by day 4. **E,** Myomaker knockout cells did not fuse but showed slight elongation following identical starvation treatment.
